# BifurcatoR: A Framework for Revealing Clinically Actionable Signal in Variance Masquerading as Noise

**DOI:** 10.1101/2025.06.01.657083

**Authors:** Zachary Madaj, Mao Ding, Carmen Khoo, Ember Tokarski, Luca Fagnocchi, PERMUTE, Ziru Li, Jesse Riordan, John Andrew Pospisilik, Joseph H. Nadeau, Christine W. Lary, Timothy J. Triche

**Affiliations:** Bioinformatics and Biostatistics Core, Van Andel Institute, Grand Rapids, MI 49503, USA; Van Andel Research Institute, Grand Rapids, MI 49503, USA; MaineHealth Institute for Research, Scarborough, ME; Roux Institute at Northeastern University, Portland, ME; Department of Anatomy and Cell Biology, Carver College of Medicine, University of Iowa, Iowa City, IA

**Keywords:** phenotypic noise, variability, epigenetics, obesity, cancer, twins, heterogeneity

## Abstract

**Background:** Disease heterogeneity is a persistent challenge in medicine, complicating both research and treatment. Standard analytical pipelines often assume patient populations are homogeneous, overlooking variance patterns that may signal biologically distinct subgroups. Variance heterogeneity (VH)—including skewness, outliers, and multimodal distributions—offers a powerful but underused lens for detecting latent etiological structures relevant to prognosis and therapeutic response.

**Methods:** A major barrier to VH analysis is the fragmented landscape of available methods, many of which rely on normality assumptions that biological data frequently violate. In addition, existing tools often require programming expertise, and clear guidance on study design considerations—such as sample size and method selection—is lacking. To address these issues, we developed BifurcatoR, an open-source software platform that simplifies the detection, modeling, and interpretation of VH. BifurcatoR integrates simulation-based method evaluation, study design recommendations, and a user-friendly web interface to support VH analysis across a range of data distributions. We benchmarked VH methods through simulation and applied BifurcatoR to two clinical datasets: acute myeloid leukemia (AML) and obesity.

**Results:** Simulation studies revealed that VH method performance is highly context-specific, varying with distribution shape, mean-variance coupling, and underlying subgroup structure. In AML, BifurcatoR identified two molecularly distinct subgroups with different treatment responses, including an EVI1-high group with significantly poorer prognosis (p < 0.005) among KMT2A-rearranged cases. In a separate study, VH analysis uncovered immunophenotypic subgroups in obesity based on gene-level discordance across monozygotic twin pairs, highlighting latent variation in adipose immune cell composition.

**Conclusions:** VH is not “noise”, biological variation without clinical relevance. Instead, VH is a structured signal that can reveal latent and clinically meaningful subtypes. BifurcatoR offers a practical, accessible framework for incorporating VH into biomedical research, with implications for biomarker discovery, patient stratification, and precision medicine.

## INTRODUCTION

Medical practice seeks to improve patient outcomes, in part by identifying patients who can benefit from targeted diagnosis, treatment, and care.^1^ This tailored approach can enhance treatment efficacy, reduce side effects, and advance our understanding of disease mechanisms. However, despite the boom in big data, where datasets can contain millions of variables, ^2-4^ uncovering meaningful patient subgroups remains a major challenge.^5,6^ The perils of subgrouping based on arbitrary thresholds, especially without compelling evidence, have been well documented.^7,8^

One statistical approach to subgroup identification is the careful examination of variance. Overdispersion, where measured variance exceeds expectation, may suggest heterogeneity in a population.^9^ Similarly, heteroscedasticity between independent groups may reflect unrecognized subgroups. These phenomena, collectively referred to as variance heterogeneity (VH), can be assessed using a range of statistical tools. While existing methods like overdispersion^9^ and heteroscedasticity testing,^10^ or more advanced techniques like mixture modeling,^11,12^ can offer insights, they are often indirect, sensitive to assumptions, and rarely incorporated into routine analysis pipelines.

Mixture models, for example, can estimate subgroup membership and distributional characteristics, but they require the user to pre-specify the number of subgroups (K), a process highly sensitive to initial assumptions. Additionally, many methods assume normality, which may obscure the very heterogeneity researchers aim to detect. Despite their potential, these VH tools remain underutilized due to their complexity and limited accessibility to non-statisticians.

To address these barriers, we developed BifurcatoR, a statistical software package designed to help identify variance heterogeneity and potential subgroups in clinical and biological datasets. BifurcatoR consists of both a simulation engine for study design and power analysis, and an analysis interface for real-world data interpretation. These capabilities are delivered through both an R package and a web-based Shiny application, making the tools accessible to users regardless of their programming expertise. We demonstrate its utility through two real-world use cases: the identification of survival-associated subgroups in acute myeloid leukemia (AML), and the detection of transcriptional immunophenotypes in obesity using monozygotic twin data. By enabling researchers to systematically explore variance and multimodality, BifurcatoR supports more nuanced and powerful clinical and biological inference.

## METHODS

BifurcatoR comprises two integrated components: a command-line R package https://github.com/VanAndelInstitute/BifurcatoR and a user-friendly Shiny web interface (https://vai-bbc.shinyapps.io/Shiny_BifurcatoR/). Both components were built using R v4.4.2. The Shiny application is hosted on the Van Andel Institute’s Shiny server, enabling easy access for users without programming expertise.

The web interface contains four modules:

**Module 1:** Power simulations for detecting differences in means and variance.

**Module 2:** Power simulations for detecting bimodality within a single group.

**Module 3:** Power simulations for comparing two bimodal distributions.

**Module 4:** Analysis interface for user-uploaded data.

Modules 1-3 allow users to define effect sizes and distributions (normal, log-normal, Weibull, and beta) and provide tools for simulating data, estimating statistical power, calculating false positive rates (FPR), and selecting between parametric and non-parametric testing options. Dispersion metrics include permuted standard deviation (perm-SD), median absolute deviation (perm-MAD), Gini’s mean difference (perm-GiniMD), all bimodality methods available in the *multimode*^13^ R package and several others detailed in the Supplementary Materials.

Module 4, the analysis portion of the software, includes wrappers for established tests (e.g., ANOVA, Levene’s test, permutation tests, Kolmogorov–Smirnov test), all the aforementioned dispersion tests, and visualizations (e.g., density plots, Cullen-Frey plots, beeswarm plots). Its design ensures reproducibility, usability, and flexibility for a wide range of research questions.

Full details of module design, permutation testing, and simulation set-up are detailed in the Supplementary Materials.

### Use Case 1: Acute myeloid leukemia

Data for the TARGET - acute myeloid leukemia (AML) analysis were obtained from *database of Genotypes and Phenotypes* (dbGaP) accessions phs000465^14^ and phs000178,^15^ and are available within BifurcatoR via ‘data(MLL)’. A WebR (https://trichelab.github.io/webR/mixtures/) was created for easy verification. Code for figure generation is available within the BifurcatoR GitHub under ‘vignettes’. Kaplan-Meier curves were generated using survfit from the *survival* package^16^ and visualized using *survminer*^*17*^. Log-rank tests assessed differences in overall survival. Bimodality was evaluated using BifurcatoR’s bs_lrt, a wrapper for *mixR*’s^11^ bs.lrt. Survival data were modeled as Weibull (non-normal) mixtures; log (gene expression) was assumed normally distributed. P-values derive from 100 bootstraps. Enrichment of EVI1 ‘high’ expression with short-term survival was tested via a chi-square test. Parameters from mixfit informed power calculations using BifurcatoR’s est_pow across sample sizes 40, 50, 60, 70, 80, 90, and 100; with 1000 simulations each. We compared the theoretical and empirical power of all bimodality methods available in BifurcatoR; empirical power was estimated by bootstrapped subsampling of the MLL data to the same respective sample sizes ranging from 40-100.

### Use Case 2: Obesity Heterogeneity

Data for the obesity-TwinsUK analysis were sourced from ArrayExpress (E-MTAB-1140) and limited to monozygotic twin pairs with data for both twins. Code is available in the BifurcatoR GitHub under ‘vignettes’. Gene expression data were restricted to the 4000 most variable genes, based on standard deviation of log-normalized count differences between cotwins. Expression deltas (Δ) were calculated as the expression in the higher-BMI twin minus that in the lower-BMI twin. This directional measure was used to identify genes potentially associated with BMI, with bifurcation expected around Δ = 0 for non-associated genes and Δ ≠ 0 for BMI-related genes.

Each gene was tested for bimodality using BifurcatoR’s bs_lrt, assuming a mixture of normally distributed data and 10,000 bootstraps. P-values were corrected via Benjamini-Hochberg to control the false discovery rate at 5%. Genes with one mode comprising <10% of samples were excluded to reduce false positives due to outliers. The resulting geneset was then compared to the UPV-B signature. ^18^ Over-representation analysis using *clusterProfiler*^*19*^ was performed on two gene sets: the bimodal genes (referred to as structural heterogeneity genes), and 65 of the 127 UPV-B genes that overlapped with the 4,000 most variable genes. Results were visualized using cnetplot from *enrichplot*.^20^

Cell type deconvolution of adipose tissue transcriptional profiles was performed using CIBERSORTx ^8^, following the approach in Yang et al.^18^ Differences in cell type proportions between twins were incorporated into a Seurat v5.1.0 pipeline with default parameters^21^ Seurat was executed with default parameters, utilizing the entire dataset for Principal Component Analysis (PCA). PCA was performed on the full dataset, and only PCs significantly associated with inter-twin cell type differences (p < 0.01, via linear regression) were used for downstream analysis.

## RESULTS

To demonstrate BifurcatoR’s utility, we outline a recommended workflow for detecting structure (e.g., subgroups) in cohort data, assess performance via simulation studies, and apply the tool to two real-world case studies: (1) AML, identifying clinically relevant subgroups via gene expression variability, and (2) obesity, revealing two inflammation-discordant endotypes.

### Recommended Workflow for Analyzing Differential Variability and Multimodal Distributions

This workflow supports both hypothesis testing and study design and is fully supported within BifurcatoR, including exploratory analysis, statistical modeling, and simulation-based validation (**Figure 1**).

**Figure 1.**
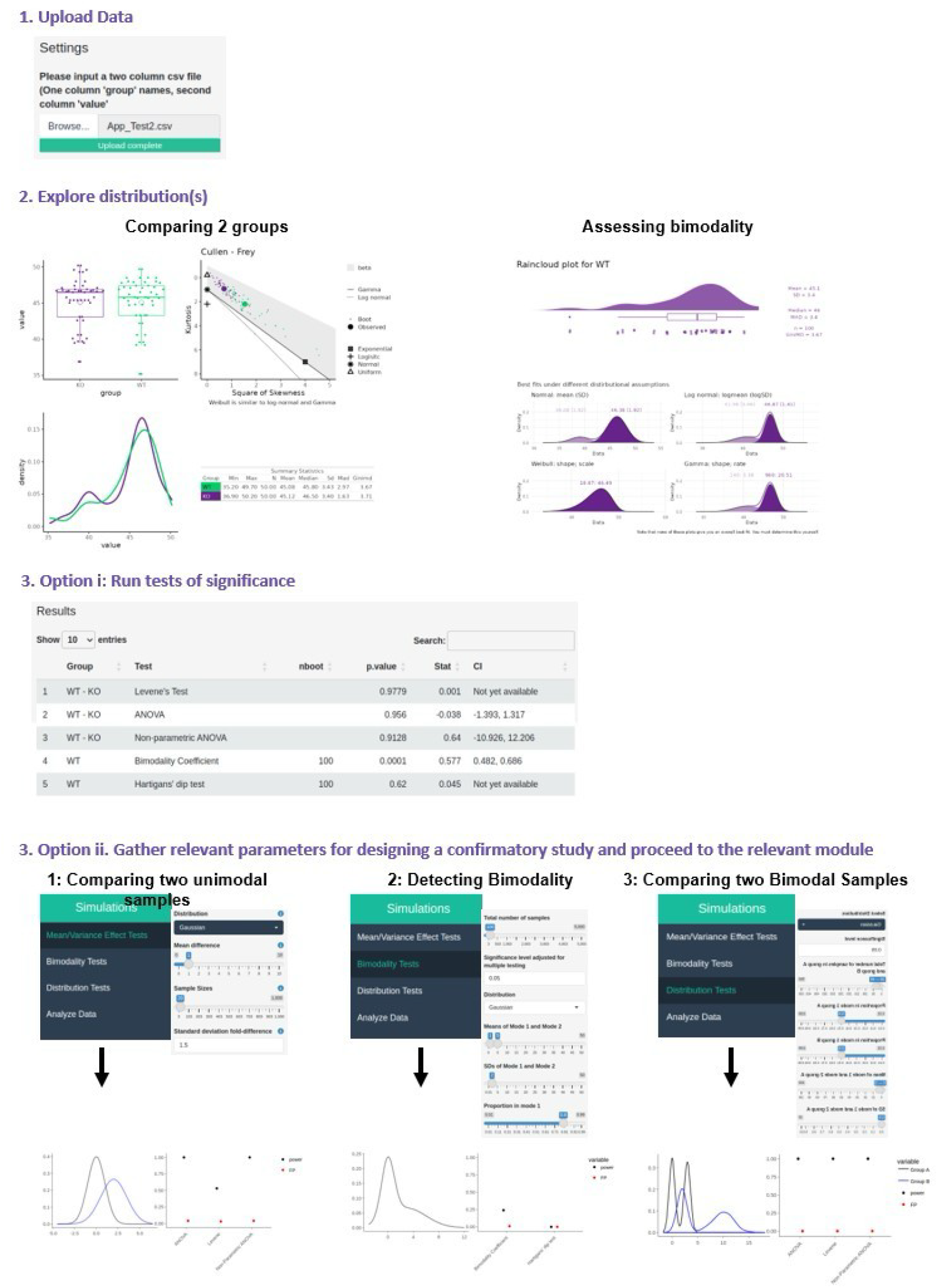
Workflow for variance heterogeneity and bimodality study planning and analysis with BifurcatoR. 1. Upload either a full dataset for analysis or pilot data 2. Perform exploratory data analysis to better understand shape, scale, and modality of data. 3. Either: i. Run desired tests of significance on a full dataset for final inference ii. Gather relevant parameters from 2. for use in a respective power analysis 1. Module 1: comparing two unimodal groups 2. Module 2: testing for bimodality 3. Module 3: comparing two bimodal groups

**Step 1**: *Data Preparation*

Users upload data comprising a categorical grouping variable (e.g., treatment and cohort) with the column header ‘group’ and a numeric outcome (e.g., gene expression and biomarker levels) value. This standardized input ensures compatibility across BifurcatoR’s analyses.

**Step 2**: *Exploratory Data Analysis*

BifurcatoR facilitates assessment of distributional properties critical for determining whether data are unimodal or multimodal. Cullen-Frey plots visualize skewness and kurtosis to guide appropriate methods: unimodal distributions (e.g., normal, log-normal or Weibull) are amenable to standard variance- and mean-based tests; multimodal distributions (e.g., beta) suggest hidden subpopulations, warranting mixture modeling, bimodality testing, or comparison of the entire distribution using methods like the Kolmogorov-Smirnov test.

**Step 3**: *Hypothesis Testing / Study Design*

The workflow diverges based on user goals:

- **Hypothesis Testing:** Users select tests such as Levene’s test (variance comparison) or Gaussian mixR (bimodality detection), in **Module 4** the **‘Analyze Data’** module, with simulation results informing test choice (expanded results are reported in the Supplementary Materials).
- **Study Design:** Users extract parameters from Step 2 (e.g., mean, SD, shape and mixing proportion) and select the appropriate power analysis module:
  ∘ **Module 1-Mean/Variance Effect Tests:** Compare two unimodal distributions
  ∘ **Module 2-Biomodality Tests:** Detect bimodality within a single dataset
  ∘ **Module 3-Distribution Tests:** Compare two multimodal datasets

Note: Users without pilot data can still use BifurcatoR for study design in this way; however, they will need to rely on educated guesses for the input parameters.

### Simulation Results: A systematic comparative evaluation of method performance

To evaluate the reliability of various statistical approaches, we performed comprehensive simulations across a range of mean and variance effects, distributions (e.g., normal, Weibull, log-normal or beta), and sample sizes; all findings are detailed in the supplementary materials. For mean differences alone, traditional methods like ANOVA and Levene’s test, demonstrated comparable power and controlled false discovery rates effectively across normal and Weibull data. When assessing variance-only effects, Levene’s test and permutation tests based on Gini or standard deviation (SD) were generally robust, though permutation (MAD) tests excelled specifically under non-normal (Weibull) distributions. Notably, combining mean and variance effects introduced complexities: in scenarios with imbalanced variance-to-sample size ratios, certain methods—especially ANOVA—showed inflated FPR, underscoring the need for caution when variance heterogeneity is present.

For bimodality detection, MixR-based methods (e.g., GmixR and WmixR) consistently outperformed alternatives in both power and specificity, particularly in identifying subtle modal differences where means overlap but variances differ (e.g., platykurtic distributions). Although the bimodality coefficient was occasionally competitive, especially for beta-distributed data (e.g., methylation β-values), it was less reliable across most other conditions. Overall, our findings demonstrate that no single method is universally optimal. Instead, test performance depends heavily on distributional characteristics and effect structure, emphasizing the importance of flexible tools like BifurcatoR for identifying patient subgroups accurately and informing clinical research design.

Finally, to validate BifurcatoR’s simulation accuracy, we compared its power and FPR estimates to those from an empirical down-sampling approach, using EVI1 (ecotropic viral integration site 1) gene expression and AML survival data from TARGET as case studies. The primary advantage of this down-sampling procedure is that it avoids making distributional assumptions; therefore, alignment between BifurcatoR’s estimates and those from the down-sampled data suggests an accurate characterization of the true underlying distribution. Results are fully detailed in the Supplementary Materials and shown in **Figure S6**. In short, there was strong concordance between both methods, confirming the reliability of BifurcatoR’s simulation module and mixR was consistently identified as a robust method for detecting bimodality in both gene expression and survival data.

### Applied data analyses and Case Studies

BifurcatoR includes two modes to support researchers: a simulation mode for selecting optimal statistical methods and ensuring adequately powered study designs, and an analytic mode for direct analysis of user-provided data to detect VH and bimodality. The platform is designed to be intuitive and accessible to researchers across disciplines, regardless of their statistical background. To illustrate its application, we analyzed two publicly available datasets: TARGET (acute myeloid leukemia) and TwinsUK (obesity); both representative of typical clinical cohorts comprising several hundred individuals and focused on complex trait diseases.

### Acute myeloid leukemia

Acute myeloid leukemia (AML) is an exceptionally heterogeneous and often lethal disease characterized by low mutational burden, diverse structural variation, and frequent resistance to combination therapies. In young (under 40) patients, a 5-year overall survival has stagnated near 68% for decades^22^.

While KMT2A rearrangements have helped define molecular subtypes, clinical outcomes remain highly variable^23^. These previous efforts to risk-stratify KMT2A-rearranged disease have largely focused on cytogenetics and fusion partner genes (examples shown in **Figure 2A**), but these markers have not reliably predicted treatment responses. To explore a gene expression–based approach, we analyzed data from 470 participants in Children’s Oncology Group AML trials, ^14,24^ investigating discordant expression of the MECOM gene (**Table 1A**).

**Table 1.** Patient demographics for the acute myeloid leukemia and monozygotic twins cohorts.

**A) Acute myeloid leukemia (TARGET; n = 470)**
| Variables | MLL Fusion* (n= 367) | NSD1 Fusion* (n = 103) |
| --- | --- | --- |
| **Age Group, n (%)** |  |  |
| Adolescent and Young Adult | 56 (15.3%) | 21 (20.4%) |
| Child | 144 (39.2%) | 76 (73.8%) |
| Infant | 166 (45.2%) | 6 (5.8%) |
| Unknown | 1 (0.3%) | 0 |
| **Sex, n (%)** |  |  |
| Male | 190 (51.7%) | 68 (66.0%) |
| Female | 176 (48.0%) | 35 (34.0%) |
| Unknown | 1 (0.3%) | 0 (0%) |
| **French-American-British classification for categorizing hematologic diseases (%)** |  |  |
| M0 | 7 (2.9%) | 3 (3.6%) |
| M1 | 14 (5.8%) | 13 (16%) |
| M2 | 10 (4.2%) | 23 (28%) |
| M4 | 105 (44%) | 24 (29%) |
| M5 | 97 (40%) | 15 (18%) |
| M6 | 0 (0%) | 2 (2.4%) |
| M7 | 7 (2.9%) | 3 (3.6%) |
| **Blast percent, n (%)** | 60.8 (28.7) | 63.6 (22.3) |
| **Mean survival (SD), years** | 3.1 (2.3) | 2.6 (2.3) |
| **Vital status, n (%)** |  |  |
| Alive | 188 (51%) | 42 (41%) |
| Deceased | 179 (49%) | 61 (59%) |

\*Fusion

| Variables | Concordant*<br>(n = 94) | Discordant A*<br>(n = 44) | Discordant B*<br>(n = 80) | Intermediate*<br>(n = 66) |
| --- | --- | --- | --- | --- |
| **Mean age (SD), years** | 61.4 (7.9) | 58.0 (11.6) | 61.6 (8.8) | 58.8 (6.2) |
| **Sex, n** |  |  |  |  |
| Female | 94 | 44 | 80 | 66 |
| **Mean BMI (SD), kg/m<sup>2</sup>** | 26.3 (4.2) | 24.9 (3.0) | 26.7 (5.2) | 27.9 (5.2) |
| **Mean height (SD), m** | 161.2 (5.9) | 159.5 (6.1) | 161.4 (5.4) | 160.0 (7.0) |
| **Weight (SD), kg** | 68.3 (11.6) | 63.5 (8.5) | 69.6 (14.1) | 71.9 (16.4) |
| **Obesity, n (%)** |  |  |  |  |
| normal weight (BMI < 25) | 77 (82%) | 42 (95%) | 61 (76%) | 45 (68%) |
| obese (BMI ≥ 25) | 13 (14%) | 2 (4.5%) | 13 (16%) | 14 (21%) |
| severely obese (BMI ≥ 30) | 4 (4.3%) | 0 (0%) | 6 (7.5%) | 7 (11%) |
| **Mean lean mass (SD), kg** | 38.9 (5.0) | 37.5 (4.5) | 39.5 (4.6) | 40.4 (7.1) |
| **Mean fat mass (SD), kg** | 27.9 (7.5) | 24.4 (5.6) | 28.5 (10.3) | 29.8 (9.8) |
| **Mean lean mass index (SD), kg/m<sup>2</sup>** | 15.0 (1.6) | 14.7 (1.4) | 15.2 (1.7) | 15.7 (2.0) |
| **Mean fat mass index (SD), kg/m<sup>2</sup>** | 10.8 (2.9) | 9.6 (2.2) | 10.9 (3.9) | 11.6 (3.5) |
| **Mean fasting serum insulin (SD), mmol/L** | 3.8 (0.6) | 3.6 (0.6) | 3.7 (0.6) | 4.0 (0.9) |

**Figure 2.**
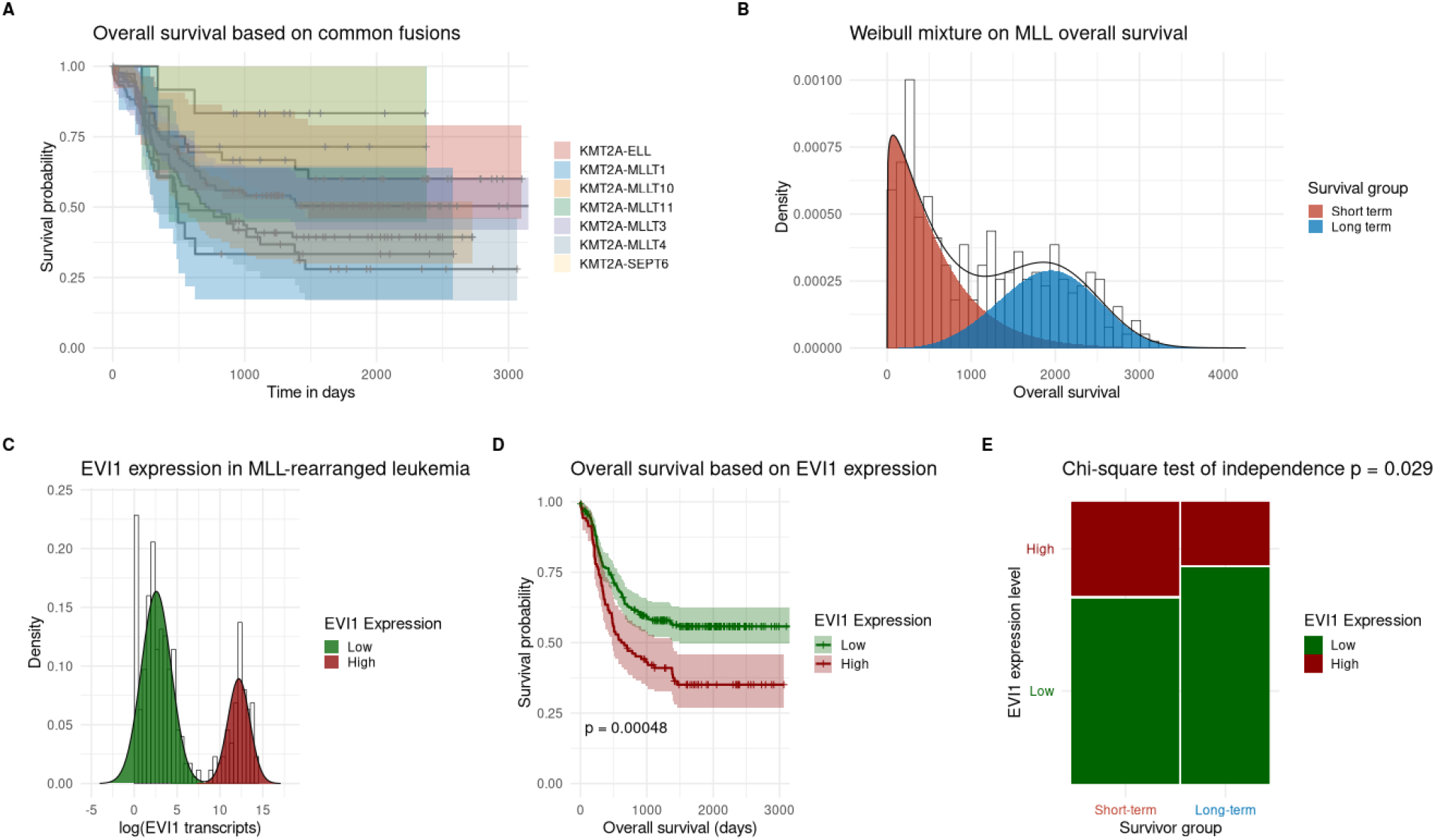
Real data analysis of within-group variation in acute myeloid leukemia. A. Kaplan-Meier curves with 95% confidence bands for fusion partners appearing in n ≥ 5 patients. B. Densities and histogram plot of overall survival generated with mixR, assuming a Weibull distribution, and revealing strong evidence for bimodality (p < 0.001). C. Densities and histogram plot of EVI1 gene expression (log2) generated with mixR using the Gaussian family, which shows strong evidence of bimodality (p < 0.001). D. Kaplan-Meier curves with 95% confidence bands for splitting the cohort into ‘low’ and ‘high’ EVI1 expression using mixR component probabilities (classification was based on the most probable mode). E. Mosaic plot of the classification matrix of EVI1 high vs low expression and long-vs short-term survival where survival was based on mixR component probabilities from B. Chi-square test on this classification table revealed significant evidence against independence between survival groups and EVI1 expression groups (p = 0.029).

Using BifurcatoR, we identified two distinct survival groups—poor (<3-year survival) and favorable (>5-year remission)—independent of fusion partner (**Figure 2B**). EVI1 (encoded by MECOM) showed strong bimodality and a 4.7-fold difference in mean expression between modes (p < 0.001; **Figure 2C**). High EVI1 expression was significantly associated with worse survival (log-rank p = 0.001; **Figure 2D**) and a chi-square test showed evidence that EVI1 expression was not independent of the survival subgroups (chi-square p = 0.029; **Figure 2E**). This case study demonstrates how BifurcatoR can uncover clinically meaningful subgroups and identify candidate biomarkers that can be linked with a simple additional chi-square test.

### Obesity subtypes

Obesity is a chronic, complex, and heterogeneous disease affecting approximately 890 million adults worldwide. Despite its strong association with comorbidities such as Type 2 diabetes, hypertension, cardiovascular disease, non-alcoholic fatty liver disease, steatohepatitis, and various malignancies,^25,26^ many individuals remain complication-free—highlighting the limitations of BMI, the primary classification tool, in capturing physiological and molecular diversity.^26-28^ This heterogeneity is believed to arise from genetic, developmental, and environmental factors that influence body composition, metabolic function, epigenetic regulation, and inflammatory profiles.^18,27-33^ The lack of precise molecular subtyping contributes to suboptimal treatment stratification, as evidenced by high nonresponse rates (10–35%) to metabolic bariatric surgery and GLP-1 receptor agonists. ^34-39^ Given obesity’s role in approximately 5 million deaths annually, addressing its biological complexity represents a critical unmet need in clinical medicine.

We analyzed gene expression discordance in 146 monozygotic twin pairs (using the leaner twin as reference), filtered to the 4,000 most variable genes, and applied BifurcatoR to test for bimodality. This yielded 292 Structured Heterogeneity (SH) genes with significant bimodal expression patterns (**Figure 3A-B**), suggesting the presence of distinct molecular subgroups. Enrichment analysis showed these SH genes were immune-related and had minimal overlap with the previously defined UPV-B (unexplained phenotypic variation – type B) gene set,^18^ which was more enriched for metabolic pathways (**Figure S7A-B**). In silico cell-type deconvolution revealed that expression differences in SH genes corresponded with divergent immune cell infiltration profiles—stratifying twin pairs into two clusters distinguished by discordant levels of adipocytes, macrophages, dendritic cells, and pericytes (**Figure 3D-E**).

**Figure 3.**
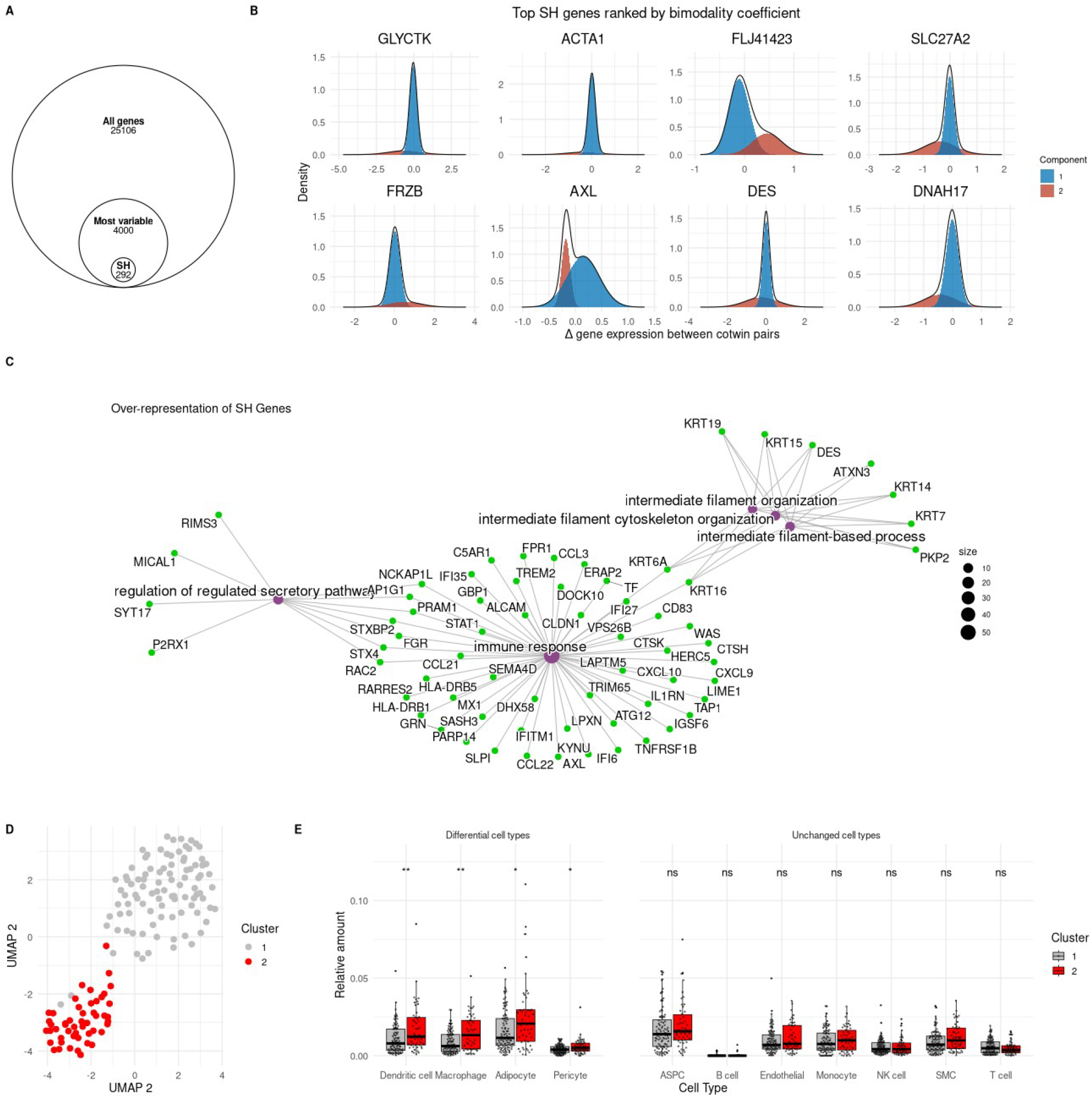
Analysis of ‘Structured Heterogeneity’ in gene expression profiles stratifies humans into two distinct metabolic state clusters with differing adipose tissue immune cell composition signatures. A. The TwinsUK data has expression measured on 25106 genes. Structured heterogeneity was investigated on the 4000 most variable genes based the gene expression discordance between cotwin pairs where 292 has significant evidence of being SH after BH multiple testing corrections. B. The densities and histograms of the top eight SH genes ranked by bimodality coefficients generated with mixR and a Gaussian family. C. Over-representation network containing gene ontologies (purple) significantly enriched with SH genes (shown in green) (FDR < 0.05). Size is the number of genes found in a given pathway. The 4000 most variable genes were used as the “background universe” D. UMAP of *in silico* estimates of cell-type proportions colored by clusters identified using Seurat E .Boxplots of each twin’s cell-type proportion split by Seurat cluster. Wilcoxon tests were used to determine if cell-type proportions differed between clusters. ‘*’ p < 0.05; “**’ p < 0.01

Importantly, SH genes did not associate with the prior UPV-B-defined obesity endotypes, indicating that BifurcatoR uncovered a distinct axis of adipose tissue heterogeneity. These findings demonstrate that even under genetically controlled conditions, adipose tissue exhibits structured immunophenotypic variation, supporting the existence of novel immuno-endotypes of obesity.

## DISCUSSION

### “Ignoring variability does not make it go away.” — Werner Kalow

Biomedical research has long prioritized mean-based analyses, often treating variability as purely statistical noise. This applies across preclinical studies, small clinical trials, and large-scale cohorts like GWAS. While productive, this approach has masked legitimate biological heterogeneity—despite longstanding recognition that the best dose for the average patient is rarely the best dose for any individual^40^. Heterogeneity in disease risk, progression, and treatment response remains underexplored due to persistent methodological barriers.^5,6^

Studying variance heterogeneity (VH) is challenging due to reliance on normality assumptions, lack of consensus, and limited accessibility. Many existing methods require statistical expertise and perform well only in specific contexts, as shown in our simulations. Even trained analysts may struggle to confidently choose the right test. **BifurcatoR** lowers this barrier incorporating study design tools and analytic workflows into one intuitive platform. Further, the expansive simulation results offer detailed guidance for choosing scenario-specific methods. Our findings show that focusing on—rather than ignoring—VH can yield rapid and clinically meaningful insights.

### Biological Insights from VH Analysis

Our case studies in AML and obesity demonstrate the value of structured VH analysis. In AML, we identified distinct patient subgroups based on the bimodality of **EVI1** expression, which strongly predicted survival across trials (log-rank *p* = 0.001; chi-square *p* = 0.029, **Figures 3D–E**). While EVI1 has been previously linked to prognosis,^41^ our findings highlight the advantages of mixture modeling over arbitrary cut-point methods like OptimalCutpoints,^42^ which are sensitive to dataset-specific noise. Mixture models define biologically grounded subpopulations, improving reproducibility across cohorts.

In the obesity dataset, we identified 292 **Structured Heterogeneity (SH)** genes via gene expression discordance in 146 monozygotic twin pairs. These genes, enriched for immune-related pathways, revealed an **immuno-inflammatory axis** of heterogeneity independent from previously defined UPV-B endotypes. ^18^ SH stratified individuals by adipose immune cell composition and suggested an environmentally driven layer of variability distinct from developmental-epigenetic effects. Given the central role of immune dysregulation in obesity-related complications,^30,33^ this analysis underscores the importance of transcriptome-based VH analysis for uncovering subtypes in genetically controlled models.

### Applications Beyond Biology

The ability to detect and interpret VH extends to regulatory agencies, IRBs, and grant reviewers, who must ensure studies are adequately powered. BifurcatoR offers a means to design such studies from the outset and gives reviewers an accessible tool to verify calculations. This may be especially crucial for clinical trials targeting precision medicine, which have stringent requirements for sample justifications.

VH methods—such as mixture modeling and structured discordance analysis—are also well established in other domains, including flow cytometry^43^ and genotype calling.^44^ In computational science, mixture models and Hidden Markov Models both support high-dimensional clustering and form the foundation of applications from speech recognition to biological sequence analysis.^45-47^ Their success in these fields points to untapped potential within biomedical research.

### Conclusion

Structured heterogeneity is not noise. It is common, biologically informative, and clinically actionable; yet often overlooked. BifurcatoR meets this challenge by providing an accessible, simulation-driven framework for VH detection and study design. As precision medicine advances, tools like BifurcatoR will be key to enabling robust biomarker discovery, patient stratification, and individualized treatment. We encourage broader adoption of VH analysis to advance understanding of disease mechanisms, environmental contributions, and therapeutic variability.

## ACKNOWLEDGEMENTS

We dedicate this work to the memory of Carmen Khoo, our friend and colleague. NIH grants AI171984, CA290259, CA251066, GM121301 and HG012444 supported this work. This work was completed in part using the Discovery cluster, supported by Northeastern University’s Research Computing team. TwinsUK data access was granted after approval of project E1106. TwinsUK is funded by the Wellcome Trust, Medical Research Council, Versus Arthritis, European Union Horizon 2020, Chronic Disease Research Foundation (CDRF), Zoe Ltd, the National Institute for Health and Care Research (NIHR) Clinical Research Network (CRN) and Biomedical Research Centre based at Guy’s and St Thomas’ NHS Foundation Trust in partnership with King’s College London.

## DATA AVAILABILITY

The experimental and clinical datasets discussed in the paper are included in the **BifurcatoR** package.

## CODE AVAILABILITY

Github repo: https://github.com/VanAndelInstitute/BifurcatoR

Shiny app link: https://vai-bbc.shinyapps.io/Shiny_BifurcatoR/

Infant leukemia vignette: https://trichelab.github.io/webR/mixtures/

TwinsUK vignette: https://trichelab.github.io/webR/twins/

Code for figures: https://github.com/VanAndelInstitute/BifurcatoRR/tree/main/vignettes

