## Supplementary Materials for "BifurcatoR: A Framework for Revealing Clinically Actionable Signal in Variance Masquerading as Noise"

### **Supplementary Context Figure**

#### **Origins of phenotypic variation and their common frequency distributions.**

A. The primary determinants of phenotypic variation are in bold, while examples of secondary contributors are displayed in regular font.
B. In the classic example, individual deviations are random about the mean (normal). Skewed data are also common in biology, specifically right-tail heavy distributions (eg fluorescence intensity, tissue volume or weight, cell density, etc). The BifurcatoR package focuses primarily on normal, bimodal, and skewed. However, specific types of platykurtic distributions (low kurtosis) are also possible to investigate; specifically, when data are characterized by a mixture of two distributions with the same mean, but different variances. Leptokurtic distributions (excessive kurtosis) are niche edge-cases outside the primary scope of the software.

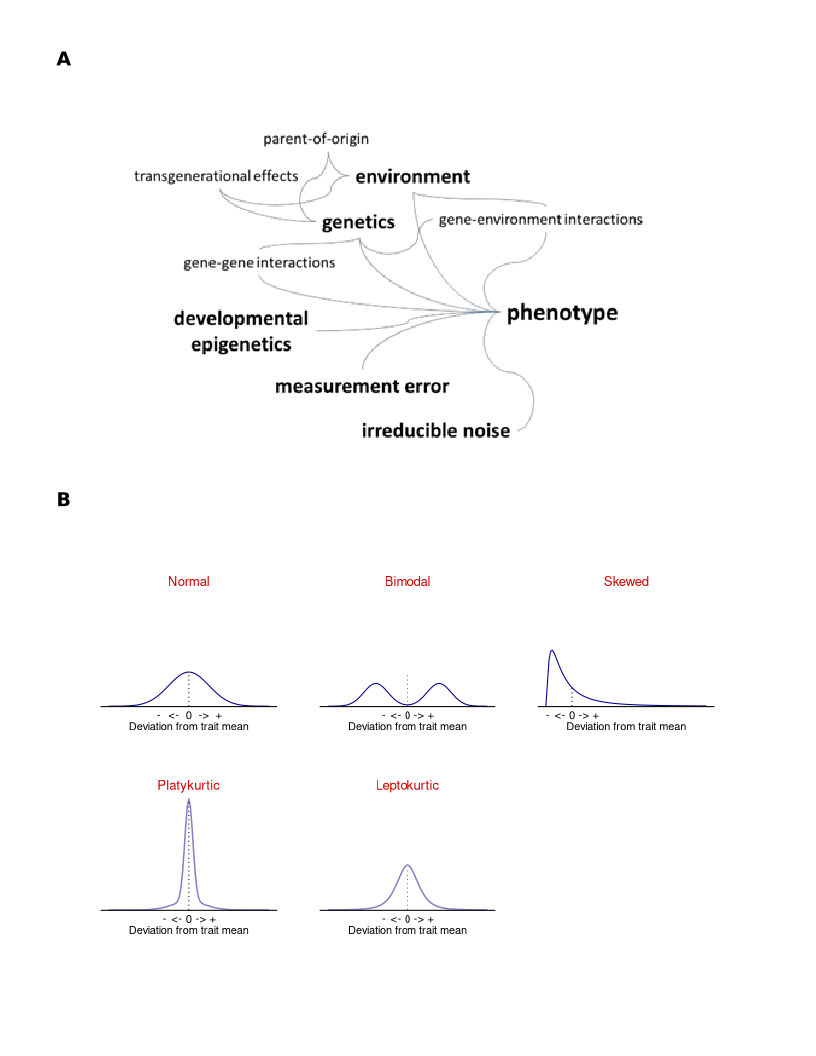

### **Supplementary Methods**

#### ***General methods***

All data were analyzed using R v 4.4.2 with visualizations made using ggplot2 v3.5.1^1^ and extensions of ggplot2 as noted in the methods. An extension of *mixR* is implemented in BifurcatoR, where a fuzzy C-means^2^ clustering is used if K-means fails; an quirk that can happen under niche situations where one cluster has ends up with a membership of n = 1.

#### ***Distributions and Effect Sizes for all modules***

All modules support normal, log-normal, and Weibull distributions, with optional beta distributions in Module 2. Beta was added to module 2 for its ability to model both unimodal and bimodal; while Weibull and log-normal are both right-skewed distributions with log-normal being particularly adept at modelling data with large variances. Users define effect sizes, and BifurcatoR automatically converts them into shape and scale parameters. *Power and False Positive Rate (FPR) Estimation:* Statistical power is calculated through repeated simulations (e.g., 1000 random datasets), and FPR is determined by generating null datasets where both groups follow the same distribution. *Measures of Dispersion:* Standard deviation (perm-SD), median absolute deviation (perm-MAD), and Gini’s mean difference (perm-GiniMD) are included to assess VH. *Parametric vs. Non-Parametric Testing:* Users can select from parametric (e.g. ANOVA) and non-parametric (e.g., permutation-based) statistical tests depending on their dataset’s characteristics.

Each module then builds upon this shared statistical framework. Critically, the Shiny application provides a user interface tailored to the distinct objectives of each type of analysis. As indicated above, the gap in knowledge in the field is the combined product of (1) a lack of tools that are ‘accessible’ to non-R users, (2) a low overall appreciation of the confounding potential and vast opportunities of multimodal analysis in the clinical space, (3) the large number of possible methodologies that can theoretically be applied to these types of analyses, and (4) importantly the wide range and context-specificity of the ultimate performance of these tools. Indeed, as the simulation analyses below highlight, some of the tools are completely powerless in select contexts. By bringing these tools all into one platform, built with biologists and non-R users in mind, BifurcatoR addresses these problems simultaneously. Each module builds on this foundation while providing additional specialized analyses. Below, we describe the specific objectives and functionality of each module in detail.

#### ***Module 1: Detecting mean and variance heterogeneity***

This module provides a framework for detecting differences in means and VH. Understanding VH is critical in clinical research, as it may indicate previously unrecognized patient subgroups (e.g., identifying differences in the extent or ‘distribution’ of treatment-response variability between two cohorts receiving the same medication).

Users can simulate datasets with normal, log-normal, and Weibull distributions by specifying a mean difference and standard deviation fold-difference between two independent groups. The reference group is standardized based on the chosen distribution model to ensure comparability. Real-time density plots allow users to validate input parameters. ***Measures of dispersion:*** Users can select from standard deviation (perm-SD), median absolute deviation (perm-MAD), and Gini’s mean difference (perm-GiniMD), which provide robust assessments of VH, particularly in skewed distributions. ***Statistical power and false positive rate (FPR) estimation:*** Power is estimated by generating a user-specified number of random datasets (e.g., 1000 simulations) and calculating the fraction that yield a significant difference at a chosen alpha level (e.g., p < 0.05). FPR is determined by simulating null datasets where both groups follow the same distribution. ***Statistical tests:*** Available methods include parametric (ANOVA, Levene’s test) and non-parametric (ranked outcome ANOVA, permutation tests) approaches. Results are presented as downloadable tables and plots.

#### ***Module 2: Detecting bimodality***

This module builds upon Module 1 by incorporating multimodal detection methods, allowing researchers to estimate power and false positive rates (FPR) for bimodal distributions and thereby enabling study design. Detecting bimodality is particularly useful in cases where a single disease diagnosis may actually encompass distinct biological subtypes (e.g., molecularly specific forms of diabetes based on biomarker distributions).

*Bimodality detection methods:* mClust^3^, Bimodality Coefficient^4^, mixR*^5^,* and from the multimode*^6^* package*:* Fisher and Marron Excess Mass test, Hall and York Bandwidth test, Hartigans’ dip test, Silverman’s Bandwidth test, Cheng and Hall excess mass test, and Ameijeiras-Alonso, Crujeiras, and Rodríguez-Casal test. S*upported distributions:* Users can simulate datasets using normal, log-normal, Weibull, or beta distributions. *Parameter specification:* In contrast to Module 1, users define parameters for each mode and specify the mixing proportion (i.e., the relative sample sizes in each mode). *Power and FPR estimation:* Real-time density plots again allow users to validate their chosen parameters. Test results are plotted and tabulated for downloading.

#### ***Module 3: Comparing two bimodal groups***

This module builds on the previous two by enabling *comparisons* between two bimodal distributions. Specifically, it incorporates methods to test whether two bimodal distributions are different (e.g. testing if two related clinical cohorts might contain different ratios of poorly understood patient subgroups, such as responders and non-responders to a targeted therapy).

***Bimodal group comparisons:*** Users specify bimodal distributions for two groups using the same parameter input structure as in Module 2. ***Mean and variance tests:*** The same statistical tests from Module 1 are available to compare group differences. ***Distribution comparison methods:*** Additional tests assess overall distributional differences using the twosamples*^7^* package, including the Kolmogorov–Smirnov test (KS), Cramer-von Mises (CVM), Decision Trees (DTS), and Anderson–Darling (AD). To avoid p-hacking or related unsound practices, the appropriate test should be selected *a priori* using Module 2, ensuring that the choice is based on the expected data characteristics rather than post hoc optimization. ***Simulation and permutation settings:*** Users define the number of simulations and permutations, with results displayed as plots and summary tables. All other functionality of Modules 1 and 2 are maintained.

#### ***Module 4: Analyzing user data***

This module provides an interactive interface for analyzing user-uploaded data, applying the statistical methods from Modules 1–3 to real-world datasets. For example, a researcher could upload patient gene expression data to test for VH, identify bimodal distributions, and compare subgroups—all within an intuitive, user-friendly interface.

***Supported analyses:*** Users can test for mean and variance differences, assess bimodality, and compare two bimodal distributions using uploaded datasets. ***Visualization tools:*** The module includes multiple plots to explore data characteristics, including 1) a beeswarm plot^8^, which groups points into a quasi-histogram with overlaid boxplots; 2) a custom Cullen-Frey plot based on the decisionSupportExtra^9^ R package; 3) density plots; and 4) a table of summary statistics (sample size, minimum, maximum, mean, median, SD, MAD, and Gini’s MD). ***Bimodality exploration:*** Additional visualization tools, including *mixR* density plots^5^, help users assess the best-fitting distribution for their data. ***Automated model selection:*** By default, BifurcatoR uses the mixR select function to determine the most appropriate modality for each distribution family.

### **Supplementary Results**

#### ***Simulation Results***

##### ***A systematic comparative evaluation of method performance***

To assess the reliability of different statistical approaches, we conducted extensive simulations across a range of mean effects, variance effects, distributions, and sample sizes. These simulations were designed to guide researchers in selecting optimal methods by identifying trade-offs in statistical power, false positive rates (FPR), and overall test performance under various conditions. A complete set of results is provided in Figures S1-S5, with a summary below. Should a researcher not have pilot data, they may instead identify the hypothesis test and the simulation scenario closest to their expected data to determine the optimal test. Additional details can be found in the BifurcatoR Github ([https://github.com/VanAndelInstitute/BifurcatoR](https://github.com/VanAndelInstitute/bifurcatoR)). These simulation results provide valuable insights for those interested in statistical methodology; importantly, a deep understanding of them is not required to effectively use BifurcatoR*.*

***Mean differences Only.*** For independent groups with equal variances, all three mean-difference methods performed nearly identically, maintaining statistical power while effectively controlling false discovery rates (~5–10%) across all sample sizes. This held true regardless of whether the data followed a normal or Weibull distribution (Figure S1).

***Variance Effects Only.*** For datasets that are normally distributed with **no mean differences**, Levene’s test, permutation (Gini) and permutation (SD) all performed similarly, while permutation tests based on the median absolute deviation (MAD) were the least powerful. Interestingly, however, when the data come from a Weibull distribution, the same permutation (MAD) test became the consistently most powerful; likely due to its robustness in non-normal data. Surprisingly, despite standard deviation being a parameter specific to normally distributed data, permutation (SD) was the second most consistently powerful method (Figure S2).

***Mean and variance Effects.*** For normally distributed data where groups had both mean and variance differences, all methods maintained sufficient power, except under conditions where one group had a much larger variance and proportionally smaller sample size (e.g., group one has a sample size 3x larger than another experimental group and group one has a standard deviation 4 larger than group 2). Under these conditions, the false positive rate increased to an unacceptable ~20%, highlighting the sensitivity of ANOVA to violations of the equal variances assumption. By contrast, when sample sizes were reversed (e.g., 1:3 ratio), statistical tests were more robust. These findings reinforce that larger variances require larger sample sizes to be effectively characterized.

For Weibull-distributed data, ranked-outcome ANOVA exhibited an extreme false positive rate (~100%) at sample sizes as small as 100, rendering it unreliable in these cases. Across all scenarios, caution is warranted when applying mean-difference tests to data with both large variance differences and non-normality.

Finally, when testing for variance effects on normally distributed data where the groups also differ in their means, Levene’s test, permutation (Gini), and permutation (SD) performed comparably; with Levene’s test being the clear favorite when mean differences are large (>4-fold). Permutation (MAD) was again the least powerful. For Weibull-distributed data, Levene’s test consistently outperformed all other methods, confirming its broad applicability.

***Bimodality Detection.*** We evaluated methods for detecting bimodality across normal, Weibull, log-normal, and beta-distributed data. MixR-based methods (GmixR for normal data, WmixR for Weibull data) consistently provided the best detection power while maintaining low false positive rates. While other methods occasionally achieved similar power, they typically had higher false positive rates, making them less reliable. A key advantage of MixR is its ability to detect bimodal distributions even when the modes have identical means but differ in variance, making it uniquely suited for identifying multiple forms of VH (e.g. platykurtic) (Figure S3, top row). The bimodality coefficient performed well when groups had equal variances, unequal sample sizes, and large mean differences (4-fold), but struggled under other conditions. Notably, for beta-distributed data, the bimodality coefficient outperformed MixR, suggesting its usefulness in data measured as a percentage that ranges from 0% to 100% such as DNA methylation β-values.

Collectively, these findings underscore the striking context dependence of method performance and the inherent challenge of identifying a ‘one-size-fits-all’ approach for detecting clinically relevant subgroups across many realistic distributions. They highlight why a tool like BifurcatoR is essential for studying patient heterogeneity and advancing clinical research.

##### ***Figure S1.* *Simulation results for mean differences on data samples from Gaussian or Weibull distributions.***

Based on 1000 simulations and 1000 permutations. Power was estimated as the fraction of times a test was called significant (p < 0.05). Parent distributions are plotted in true red (reference group) and true blue above each panel. The left column of each set of 9 plots shows results for mean-only differences; while the top row is variance-only effects. Notably, for simulations where the two independent groups have no mean difference, some power estimates are as high as 0.99. For these calculations, the tests are correctly rejecting the null model (specifically, the equal variances assumption or exchangeability assumption). However in practice, these results may be falsely interpreted as the null hypothesis of no mean difference; demonstrating the importance of validating model assumptions prior to making inferences.

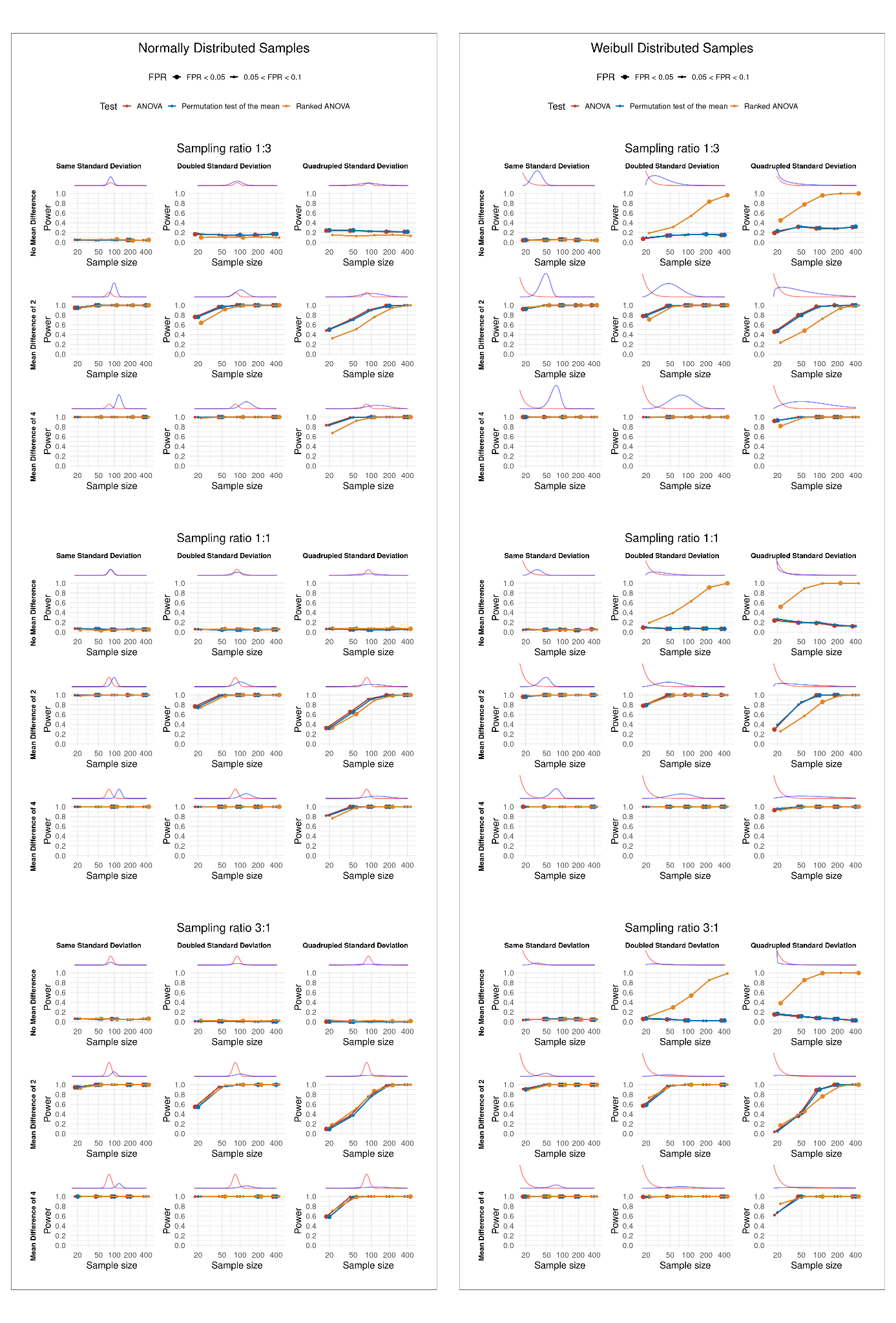

##### ***Figure S2. Simulation results for variance methods (Module 1)***

Simulations (n=1000) were run for sample sizes 20, 50,100, 200, and 400 with the goal of estimating the power to detect heteroscedasticity between two independent samples and simultaneously estimate the FPR for several variance tests. Data were drawn from either normal or Weibull distributions with various mean and variance differences between the two groups (no mean difference, 2-fold difference in means, 4-fold difference in means, and with the same differences for standard deviations). The proportion of samples drawn from each group was also varied, a 1:3 sampling ratio, even split, and 3:1. Parent distributions are plotted in true blue and true red above each panel, with group 1 (reference group) in red and group 2 (test group) in blue. FPRs were binned into FPR < 0.05; 0.05 < FPR < 0.1; and FPR > 0.1. Permutation methods used 1000 resamples. *MAD:* median absolute deviation; *Gini:* Gini’s mean difference; *SD:* standard deviation.
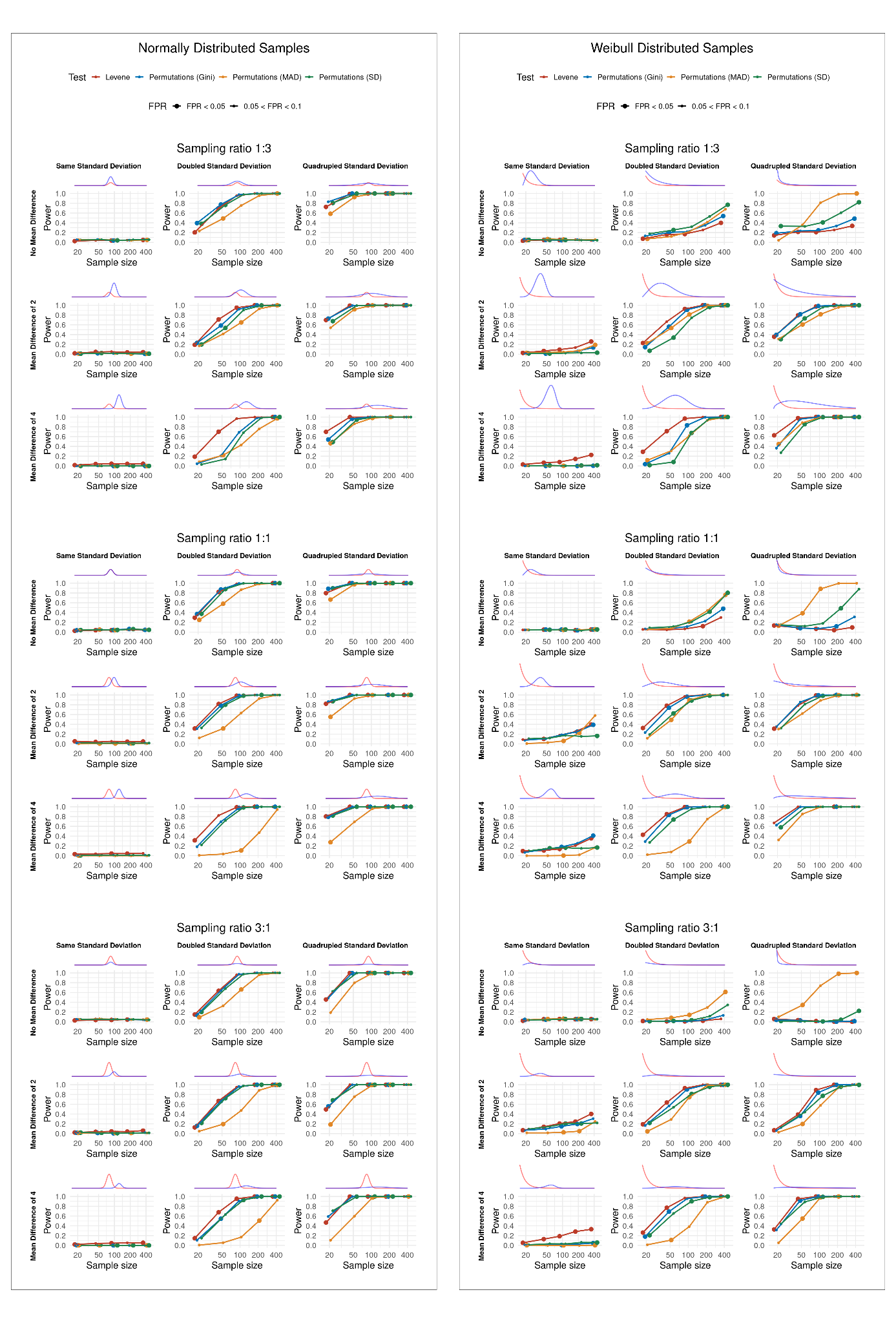

##### ***Figure S3. Simulation results for bimodality methods (Module 2)***

Simulations (n=1000) were run for sample sizes 20, 50,100, 200, and 400 with the intended goal of estimating the power to detect bimodality in a single sample and simultaneously estimate the FPR across many multimodality tests. Data were drawn from either a mixture of normal distributions or a mixture of Weibull distributions with various mean and variance differences between the two modes (no mean difference, 2-fold difference in means, 4-fold difference in means, and with the same differences for standard deviations). The proportion of samples drawn from each mode also varied, a 1:3 sampling ratio, even split, and 3:1. Parent distributions are plotted in true black above each panel. FPRs were binned into FPR < 0.05; 0.05 < FPR < 0.1; and FPR > 0.1. Permutation methods used 1000 resamples.. *MAD:* median absolute deviation; *Gini:* Gini’s mean difference; *SD:* standard deviation.

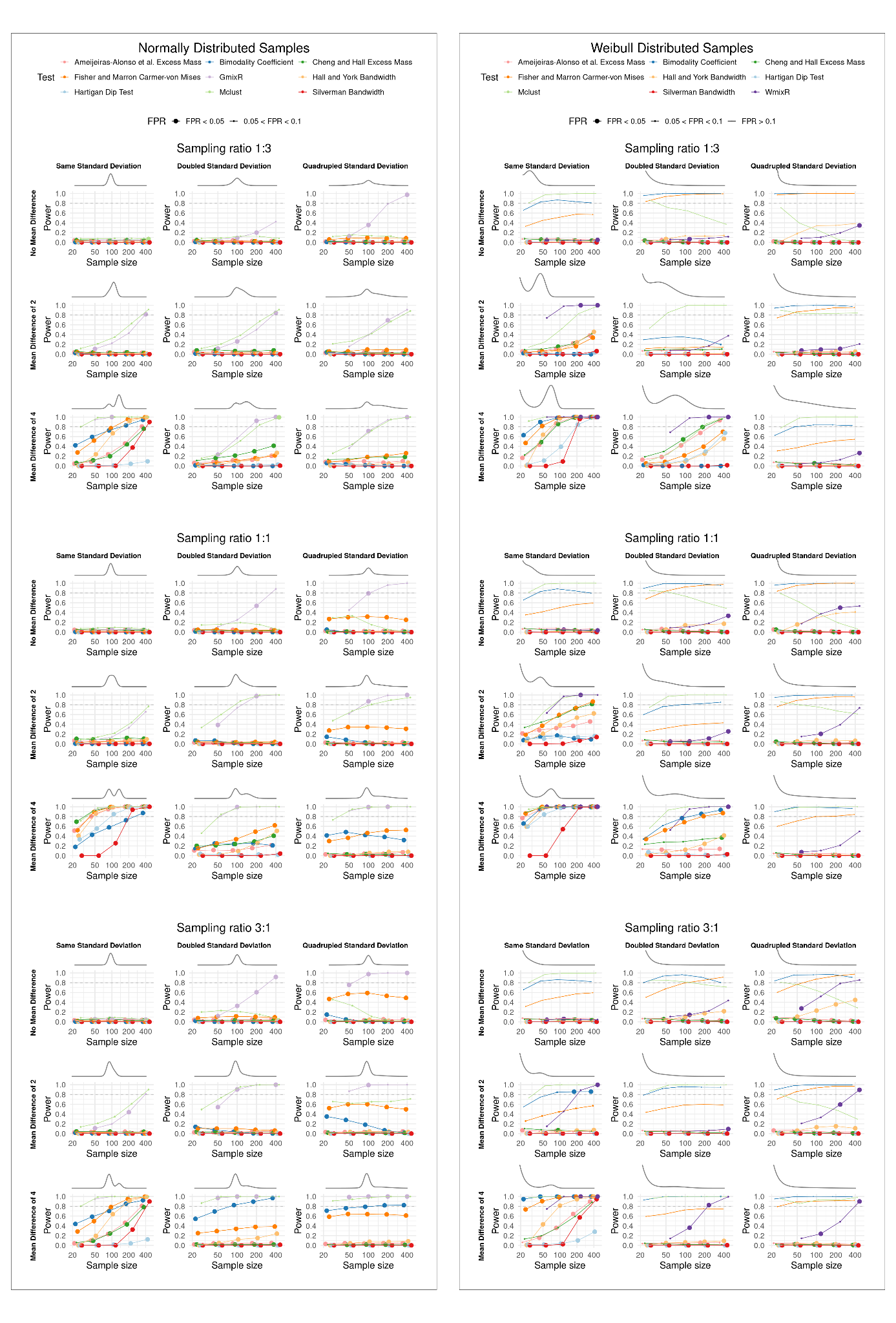

##### ***Figure S4. Simulation results for bimodality detection using log-normal mixR***

Based on 1000 simulations and 1000 permutations. Power was estimated as the fraction of times a test was called significant (p < 0.05). Parent distributions are plotted in true black above each panel.

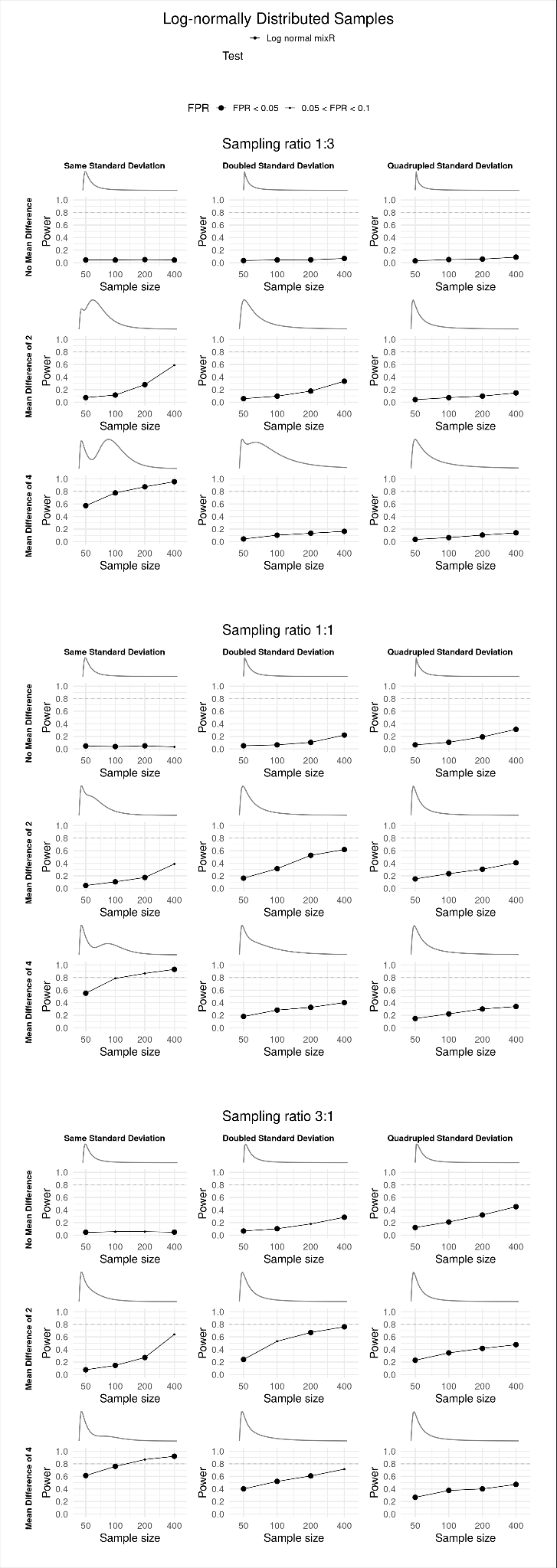

##### ***Figure S5. Simulation results for bimodality detection when data are beta-distributed***

Based on 1000 simulations and 1000 permutations. Power was estimated as the fraction of times a test was called significant (p < 0.05). Parent distributions are plotted in true black above each panel.

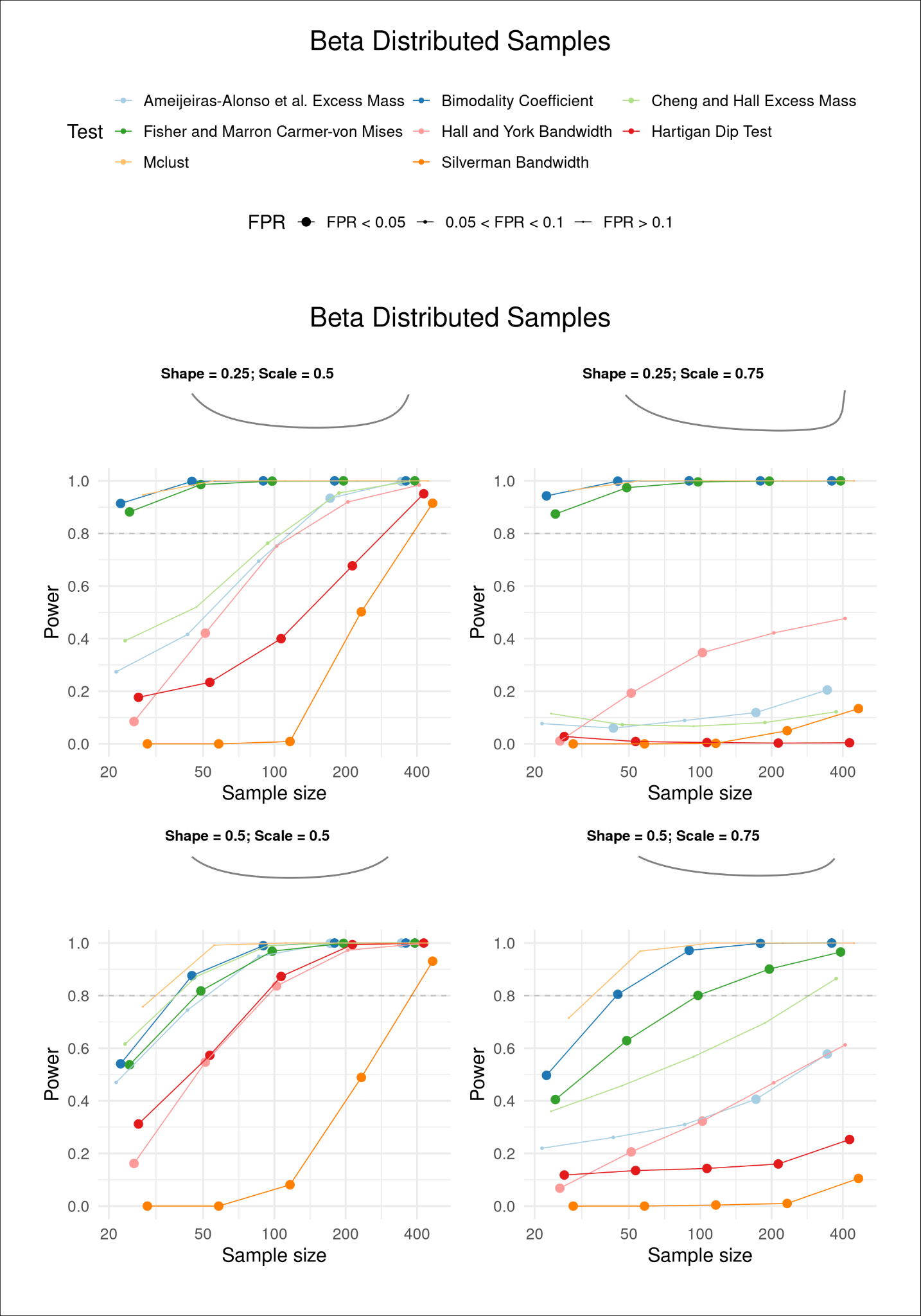

##### ***Empirical versus Theoretical Power***

We performed simulation studies to validate BifrucatoR’s simulation accuracy and thus its suitability to inform sample size and method choice for other studies looking to test genes similar to EVI1 (Figure S6A) or overall survival (Figure S6B). These simulations used the methods in BifurcatoR as well as an empirical down-sampling procedure to validate the accuracy of BifurcatoR’s simulated estimates. The down-sampling procedure creates 1000 bootstrap samples of size N and the proportion of significant tests is the ‘empirical power’ for a given method. The down-sampling method’s advantage is that it does not make any distributional assumptions, therefore for the down-sampling and BifurcatoR simulations to match, the parameter estimation within BifurcatoR must be accurate. The two sets of calculations yielded markedly consistent results, affirming BifurcatoR’s simulation modules are producing accurate power and FPR estimates. These findings also reaffirmed mixR as an optimal method when high power (>80%) is considered in conjunction with a low FPR (< 5%). Interestingly, for EVI1 expression, several other tests also showed strong detection performance (power > 0.8 and FPR < 0.05): Mclust, bimodality coefficient, and the Cheng and Hall excess mass test. For overall survival, sufficient power was not achieved until around n = 70 with Weibull mixR. Among bimodality tests considered, Hartigan’s dip test and Silverman bandwidth tests were consistently the least powerful. Putting these findings together, when looking for bimodality in expression data, there are a few viable methods with mixR coming out slightly ahead and requires as few as 40 samples; while mixR is the clear preference for time-to-event data and requires nearly twice as many samples.

##### ***Figure S6. Power and False Positive Rates under Empirical and Theoretical simulations***

A. *Left*: the estimated power and FDR range based on 1000 simulations run on sample sizes ranging from 40-100 using bimodal parameters estimated with Gaussian mixR on the EVI1 expression data: mu_1_ = 2.6, mu_2_ =12.2, SD_1_ = 1.7, SD_2_ = 1.3, proportion in EVI1 low = 0.71. *Right*: the empirical power of various sample sizes estimated by randomly down-sampling the cohort 1000 times for each N and running each subsample through BifuiractoR’s bs_lrt test. This plot does not contain FDR as a simulated null distribution is not created with down-sampling.
B. *Left*: the estimated power and FDR range based on 1000 simulations run on sample sizes ranging from 40-100 using bimodal parameters estimated with Weibull mixR on the overall survival (OS) data: scale_1_ = 1.1, scale_2_ =3.8, shape_1_ = 583, shape_2_ = 2112 , proportion in short-term survival = 0.58. *Right*: the empirical power of various sample size estimated by randomly down-sampling the cohort 1000 times for each N and running each subsample through BifruactoR’s bs_lrt test; this plot does not contain FDR as a simulated null distribution is not created with down-sampling.

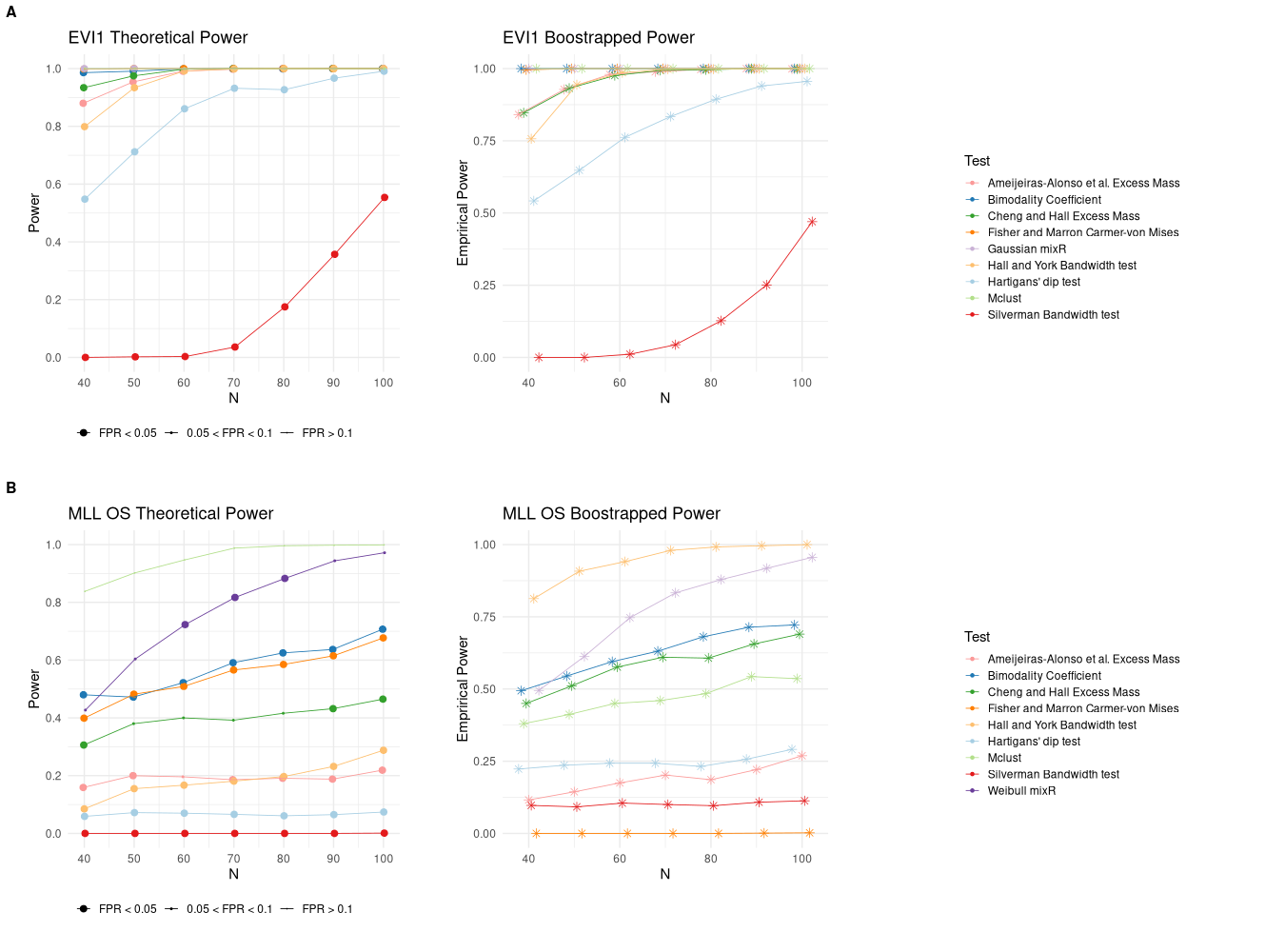

#### ***UK Twins***

##### ***Figure S7. Comparing SH genes and cell-type composition to UPV-B genes and cotwin clusters***

A) Overlap between genes identified as differentially dispersed and those found in the UPV-B signature^10^. B) Over-representation network containing gene ontologies (purple) significantly enriched with the 65 UPV-B genes (shown in green) present in the 4000 most variable genes (FDR < 0.05). Size is the number of genes found in a given pathway. The 4000 most variable genes were used as the “background universe”. C) Heatmap of z-scroed *in silico* cell-type estimates for each twin (rows). D) Percentage of cotwin pairs in Seurat clusters 1 or 2, split by previously identified cotwin clusters^10^.
B)
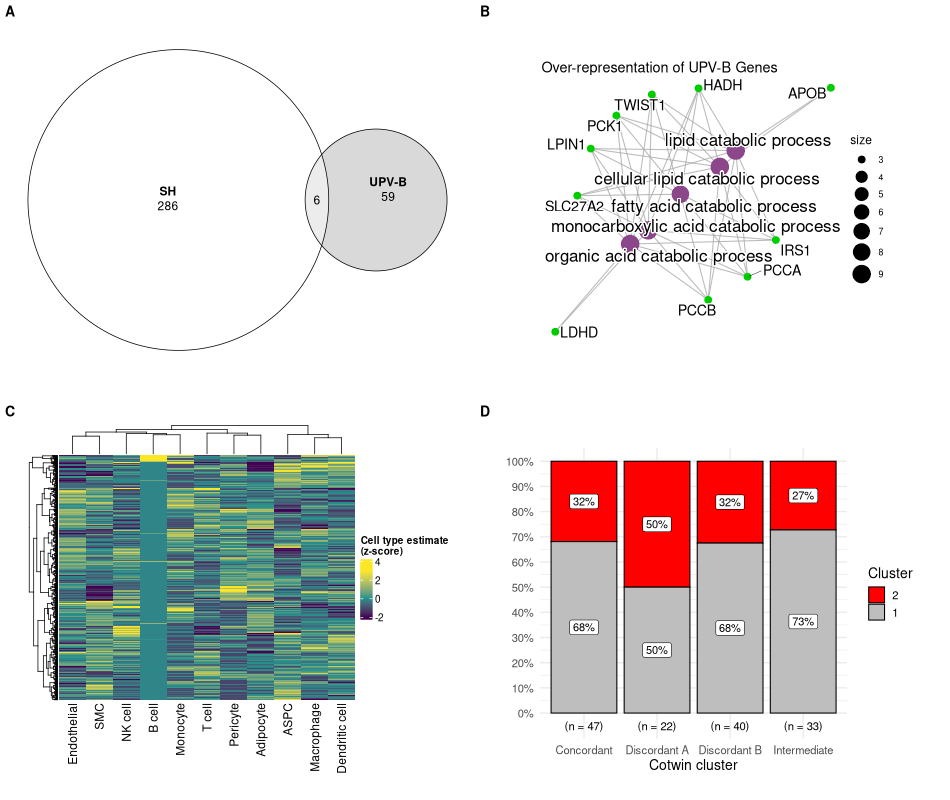

##### ***Table S1. All SH (Structured Heterogeneity) genes in the MuTHER/TwinsUK cohort.***

| **Gene** | **FDR** |
| --- | --- |
| **AACS** | **0.0248** |
| **AADACL1** | **<0.001** |
| **ACTA1** | **<0.001** |
| **ALCAM** | **<0.001** |
| **ALDH2** | **0.0058** |
| **AMOT** | **0.0265** |
| **ANKRD33** | **0.0087** |
| **ANKRD38** | **0.0369** |
| **AP1G1** | **0.0343** |
| **AQP7P2** | **0.0087** |
| **AQP9** | **0.0024** |
| **ARHGEF10** | **0.0493** |
| **ATG12** | **0.0258** |
| **ATP6AP1** | **0.0141** |
| **ATP6V1B1** | **<0.001** |
| **ATP6V1B2** | **0.0328** |
| **ATXN3** | **0.0073** |
| **AXL** | **0.0175** |
| **BTF3** | **0.0328** |
| **C10orf125** | **0.0175** |
| **C12orf39** | **0.032** |
| **C19orf28** | **<0.001** |
| **C1orf162** | **0.0428** |
| **C5AR1** | **0.0452** |
| **C5orf23** | **0.0225** |
| **CA3** | **0.0024** |
| **CBS** | **0.0042** |
| **CCDC3** | **0.031** |
| **CCL21** | **0.0087** |
| **CCL22** | **0.0024** |
| **CCL3** | **<0.001** |
| **CCNB2** | **<0.001** |
| **CCND2** | **0.0024** |
| **CCNF** | **0.0248** |
| **CD300LB** | **<0.001** |
| **CD72** | **0.0405** |
| **CD83** | **0.0214** |
| **CDCA5** | **0.0024** |
| **CEACAM6** | **<0.001** |
| **CHRDL1** | **0.0272** |
| **CHST13** | **0.0042** |
| **CHST3** | **0.0214** |
| **CISH** | **<0.001** |
| **CLDN1** | **0.0024** |
| **CLIC5** | **0.0154** |
| **CLIC6** | **0.0238** |
| **CLN3** | **0.0199** |
| **CNN1** | **<0.001** |
| **COMP** | **<0.001** |
| **COX6BP1** | **0.0214** |
| **CPA4** | **0.0248** |
| **CRTAC1** | **<0.001** |
| **CRYAB** | **0.013** |
| **CSF2RA** | **0.0154** |
| **CTSB** | **0.0351** |
| **CTSH** | **0.0073** |
| **CTSK** | **0.0087** |
| **CXCL10** | **0.0073** |
| **CXCL9** | **0.0258** |
| **CYP1A1** | **<0.001** |
| **CYP27A1** | **0.0175** |
| **DES** | **<0.001** |
| **DHX58** | **<0.001** |
| **DNAH17** | **<0.001** |
| **DOCK10** | **0.0265** |
| **DPEP2** | **0.0042** |
| **DPT** | **0.0102** |
| **DTNA** | **0.0058** |
| **E2F2** | **<0.001** |
| **EBI2** | **0.0042** |
| **ECGF1** | **0.0399** |
| **EDG4** | **0.0396** |
| **EEF1A1** | **<0.001** |
| **EGR2** | **<0.001** |
| **ELOVL6** | **0.0024** |
| **EMR2** | **<0.001** |
| **ENDOGL1** | **0.0452** |
| **EPHB6** | **0.0141** |
| **ERAP2** | **<0.001** |
| **FAM46C** | **0.0265** |
| **FAP** | **0.03** |
| **FBLN5** | **0.0387** |
| **FBP1** | **0.0087** |
| **FCF1** | **0.0073** |
| **FERMT3** | **0.0024** |
| **FGR** | **0.0024** |
| **FHOD3** | **0.0058** |
| **FKBP5** | **0.0199** |
| **FLJ13305** | **0.0024** |
| **FLJ41423** | **0.0042** |
| **FLNC** | **0.0272** |
| **FOSB** | **0.0141** |
| **FPR1** | **0.03** |
| **FRZB** | **<0.001** |
| **GABARAP** | **0.0281** |
| **GAL** | **<0.001** |
| **GBP1** | **<0.001** |
| **GDF15** | **0.0042** |
| **GLYAT** | **0.0175** |
| **GLYCTK** | **<0.001** |
| **GRN** | **0.0343** |
| **GSTM1** | **<0.001** |
| **GSTM2** | **0.0265** |
| **GSTO1** | **<0.001** |
| **HBD** | **0.0058** |
| **HBQ1** | **0.0186** |
| **HERC5** | **<0.001** |
| **HEXB** | **0.0058** |
| **HK3** | **0.0024** |
| **HLA-DRB1** | **<0.001** |
| **HLA-DRB5** | **<0.001** |
| **HP** | **0.0165** |
| **HS3ST2** | **0.0141** |
| **IFI27** | **0.0207** |
| **IFI30** | **0.0024** |
| **IFI35** | **<0.001** |
| **IFI6** | **<0.001** |
| **IFITM1** | **0.0087** |
| **IGSF6** | **0.0024** |
| **IKZF1** | **<0.001** |
| **IL1RN** | **0.0141** |
| **INMT** | **0.0281** |
| **IRX2** | **0.0465** |
| **ITLN1** | **<0.001** |
| **KRBA2** | **0.0042** |
| **KRT14** | **0.0024** |
| **KRT15** | **<0.001** |
| **KRT16** | **<0.001** |
| **KRT19** | **<0.001** |
| **KRT6A** | **<0.001** |
| **KRT7** | **<0.001** |
| **KYNU** | **0.0087** |
| **LAPTM5** | **<0.001** |
| **LDHD** | **0.0175** |
| **LIME1** | **<0.001** |
| **LOC221136** | **<0.001** |
| **LOC283050** | **0.0024** |
| **LOC285095** | **0.0024** |
| **LOC401206** | **0.0024** |
| **LOC440503** | **0.0207** |
| **LOC441763** | **0.0024** |
| **LOC642113** | **0.0024** |
| **LOC642118** | **0.0214** |
| **LOC642299** | **0.0225** |
| **LOC643318** | **0.0024** |
| **LOC644366** | **<0.001** |
| **LOC647450** | **0.013** |
| **LOC649088** | **<0.001** |
| **LOC649150** | **0.0481** |
| **LOC649923** | **0.0073** |
| **LOC649970** | **<0.001** |
| **LOC652493** | **0.0042** |
| **LOC728715** | **0.0042** |
| **LOC731486** | **0.0369** |
| **LPXN** | **0.0042** |
| **LRFN5** | **0.0472** |
| **LY6E** | **<0.001** |
| **MACROD1** | **0.0165** |
| **MAFA** | **<0.001** |
| **MAMDC4** | **<0.001** |
| **MAPK13** | **<0.001** |
| **MAT2A** | **0.0405** |
| **MATK** | **<0.001** |
| **MBTD1** | **0.0073** |
| **MED16** | **0.0073** |
| **MFN2** | **<0.001** |
| **MGAT1** | **0.0207** |
| **MGC33556** | **<0.001** |
| **MICAL1** | **0.0415** |
| **MID1IP1** | **0.0265** |
| **MMP7** | **0.0058** |
| **MOGAT1** | **0.0024** |
| **MOSC1** | **0.0165** |
| **MPP1** | **0.0186** |
| **MRPS15** | **0.0405** |
| **MRPS18A** | **0.0165** |
| **MS4A6E** | **<0.001** |
| **MT1F** | **0.0058** |
| **MT1G** | **<0.001** |
| **MUSTN1** | **<0.001** |
| **MVD** | **0.0024** |
| **MX1** | **<0.001** |
| **MYO9B** | **0.0499** |
| **MYOZ1** | **0.0024** |
| **NCKAP1L** | **0.0024** |
| **NPC1L1** | **<0.001** |
| **NPL** | **0.0024** |
| **NRIP3** | **<0.001** |
| **NXNL1** | **<0.001** |
| **OLR1** | **<0.001** |
| **OSCAR** | **0.0024** |
| **P2RX1** | **<0.001** |
| **P2RY11** | **0.0493** |
| **PAFAH1B3** | **0.0087** |
| **PAQR4** | **0.0042** |
| **PARP12** | **0.0207** |
| **PARP14** | **<0.001** |
| **PCDH17** | **<0.001** |
| **PDE1B** | **0.03** |
| **PDK2** | **0.0058** |
| **PENK** | **0.0281** |
| **PGM1** | **0.0265** |
| **PIGX** | **0.0272** |
| **PKD1L2** | **<0.001** |
| **PKP2** | **<0.001** |
| **PLA2G7** | **0.0465** |
| **PLAC9** | **0.0488** |
| **PLCB2** | **<0.001** |
| **PLIN** | **0.0387** |
| **PODXL2** | **0.0281** |
| **PRAM1** | **<0.001** |
| **PRND** | **<0.001** |
| **PRODH** | **<0.001** |
| **PRPH** | **0.013** |
| **PRR4** | **<0.001** |
| **PTTG1** | **<0.001** |
| **RAC2** | **0.0428** |
| **RARRES1** | **0.0058** |
| **RARRES2** | **0.0042** |
| **RASL11B** | **0.0042** |
| **RASL12** | **0.0141** |
| **RASSF7** | **0.0175** |
| **RBP4** | **0.0458** |
| **RDBP** | **0.0362** |
| **RGS17** | **0.0024** |
| **RHPN2** | **0.0175** |
| **RIMS3** | **0.0087** |
| **RNPEP** | **<0.001** |
| **RPS23** | **0.0396** |
| **RUNDC2C** | **0.0265** |
| **S100A3** | **0.0141** |
| **S100P** | **<0.001** |
| **SASH3** | **0.0102** |
| **SCGB1D2** | **0.0024** |
| **SCGB2A1** | **<0.001** |
| **SCNN1D** | **0.0058** |
| **SDS** | **<0.001** |
| **SEMA4D** | **<0.001** |
| **SFRS2** | **0.0336** |
| **SH3BGRL3** | **0.0024** |
| **SLC16A10** | **<0.001** |
| **SLC16A3** | **0.0042** |
| **SLC25A10** | **0.0378** |
| **SLC25A34** | **0.031** |
| **SLC27A2** | **<0.001** |
| **SLC29A3** | **0.0102** |
| **SLC35C1** | **0.0405** |
| **SLC7A1** | **0.0154** |
| **SLPI** | **0.0258** |
| **SPC24** | **<0.001** |
| **SPOCD1** | **<0.001** |
| **SQRDL** | **0.0399** |
| **SRL** | **0.0186** |
| **SRXN1** | **0.0493** |
| **ST14** | **<0.001** |
| **ST3GAL2** | **0.0165** |
| **STAT1** | **<0.001** |
| **STMN2** | **0.0024** |
| **STX4** | **0.0186** |
| **STXBP2** | **0.0087** |
| **SYT17** | **<0.001** |
| **TAF13** | **0.0058** |
| **TAP1** | **<0.001** |
| **TBL3** | **0.013** |
| **TELO2** | **0.0458** |
| **TF** | **0.0428** |
| **TFPT** | **0.0207** |
| **TK1** | **0.0024** |
| **TM7SF4** | **0.0024** |
| **TNC** | **0.0042** |
| **TNFRSF1B** | **0.0058** |
| **TNS1** | **0.0351** |
| **TNS3** | **0.0369** |
| **TOP2A** | **<0.001** |
| **TREM2** | **<0.001** |
| **TRIM65** | **0.0472** |
| **TRMU** | **0.0399** |
| **TTYH3** | **0.0058** |
| **TUBB2B** | **0.0258** |
| **UBXD2** | **0.0272** |
| **USP34** | **0.0488** |
| **VENTX** | **0.0073** |
| **VMO1** | **0.0042** |
| **VPS26B** | **0.0058** |
| **VWCE** | **0.0214** |
| **WAS** | **0.0272** |
| **WDR86** | **0.0343** |
| **XAF1** | **<0.001** |
| **ZCCHC14** | **0.0087** |
| **ZDHHC24** | **0.0058** |
| **ZFAND5** | **0.0073** |
| **ZRANB1** | **0.0024** |

### **References**

1.	Wickham H, Chang W, Henry L, et al. ggplot2: Create Elegant Data Visualisations Using the Grammar of Graph ics.

2.	Cebeci Z, Yildiz F, Kavlak AT, Cebeci C, Onder H. ppclust: Probabilistic and Possibilistic Cluster Analysis.

3.	Scrucca L, Fop M, Murphy TB, Raftery AE. mclust 5: Clustering, Classification and Density Estimation Using Gaussian Finite Mixture Models. R J 2016;8(1):289-317. (<https://www.ncbi.nlm.nih.gov/pubmed/27818791>).

4.	Kieslich PJ, Wulff DU, Henninger F, Haslbeck JMB. mousetrap: Process and Analyze Mouse-Tracking Data.

5.	Yu Y. mixR: An R package for Finite Mixture Modeling for Both Raw and Binned Data. Journal of Open Source Software 2022;7(69). DOI: 10.21105/joss.04031.

6.	Ameijeiras-Alonso J, Crujeiras RM, Rodriguez-Casal A. multimode: An R Package for Mode Assessment. Journal of Statistical Software 2021;97(9). DOI: 10.18637/jss.v097.i09.

7.	Dowd C. twosamples: Fast Permutation Based Two Sample Tests.

8.	Clarke E, Sherrill-Mix S, Dawson C. ggbeeswarm: Categorical Scatter (Violin Point) Plots.

9.	Liman I. decisionSupportExtra: Miscellaneous Functions for Quantitative Support of Decision Making under Uncertainty.

10.	Yang CH, Fagnocchi L, Apostle S, et al. Independent phenotypic plasticity axes define distinct obesity sub-types. Nat Metab 2022;4(9):1150-1165. DOI: 10.1038/s42255-022-00629-2.
